# Automated generation of context-specific Gene Regulatory Networks with a weighted approach in D. melanogaster

**DOI:** 10.1101/2020.08.03.232959

**Authors:** Leandro Murgas, Sebastian Contreras-Riquelme, J. Eduardo Martínez, Camilo Villaman, Rodrigo Santibáñez, Alberto J.M. Martin

**Affiliations:** Centro de Genónica y Bioinformática, Facultad de Ciencias, Universidad Mayor, Santiago, 8580745, Chile; Facultad de Ciencias de la Vida, Universidad Andres Bello, Santiago, 8370146, Chile; Centro de Modelamiento Molecular, Biofísica y Bioinformática – CM2B2, Facultad de Ciencias Químicas y Farmaceuticas, Universidad de Chile, Santiago, 8380492, Chile

**Author notes:** both authors contributed equally (Leandro Murgas, Sebastian Contreras-Riquelme).

**Keywords:** Systems biology, Gene regulation, Data integration, Condition-specific networks, Cytoscape

## Abstract

**Motivation:** The regulation of gene expression is a key factor in the development and maintenance of life in all organisms. This process is carried out mainly through the action of transcription factors (TFs), although other actors such as ncRNAs are involved. In this work, we propose a new method to construct Gene Regulatory Networks (GRNs) depicting regulatory events in a certain context for *Drosophila melanogaster*. Our approach is based on known relationships between epigenetics and the activity of transcription factors.

**Results:** We developed method, Tool for Weighted Epigenomic Networks in *D. melanogaster* (Fly T-WEoN), which generates GRNs starting from a reference network that contains all known gene regulations in the fly. Regulations that are unlikely taking place are removed by applying a series of knowledge-based filters. Each of these filters is implemented as an independent module that considers a type of experimental evidence, including DNA methylation, chromatin accessibility, histone modifications, and gene expression. Fly T-WEoN is based on heuristic rules that reflect current knowledge on gene regulation in *D. melanogaster* obtained from literature. Experimental data files can be generated with several standard procedures and used solely when and if available.Fly T-WEoN is available as a Cytoscape application that permits integration with other tools, and facilitates downstream network analysis. In this work, we first demonstrate the reliability of our method to then provide a relevant application case of our tool: early development of *D. melanogaster*.

**Availability:** Fly T-WEoN, together with its step-by-step guide is available at https://weon.readthedocs.io

## 1 Introduction

The regulation of gene expression is indispensable for adaptation to ever changing contexts and every aspect involved in sustaining life. Gene regulation is mainly carried out by highly specialized proteins, among which, Transcription Factors (TFs) are generally accepted as the key actors [22]. Canonically speaking, the regulation of gene expression works through the binding of TFs to certain sites in the chromatin, TF Binding Sites (TFBSs), and TFs recognize specific DNA patterns called TF Binding Motifs (TFBMs). These sites are usually unique for each TF, and they are commonly located around the promoter of TF-target genes upstream of their transcription start site. Whereas proximal upstream location of TFBS are easily related to the regulation of specific genes [25, 17], to determine which genes are controlled by each TF binding to enhancer regions have shown a greater difficulty [16, 21, 40]. Moreover, gene expression can be defined as the process by which the final products encoded by genes are generated, and thus, their regulation can also include control of translation and RNA degradation. In this way, several other non-TF regulatory elements are involved in the regulation of gene expression. For example, miRNAs and other ncRNAs are known to act during translation by binding to other RNAs [8, 3], while histone modifiers attach or remove post-translational modifications to control the positions of the chromatin that are available to be occupied by TFs.

Several epigenetic marks, including histone modifications [4] and DNA methylation [30], have been related to active and inactive states of chromatin [5, 15], therefore, influencing the ability of TFs to regulate gene expression. In this way, combinations of epigenetic marks have been related to a specific effect on TF-binding and gene expression, coining an epigenetic code that is still not properly understood [4, 2]. Even so, there are some generally accepted facts on the relationship between TF-binding and epigenetic marks that have made possible to grasp a general tendency [33]. Nonetheless, chromatin structure and epigenetic marks change dynamically in a context-specific manner, and those changes have been subject of both static and dynamic modeling to predict gene expression [36].

Despite the relationship between epigenetic marks and gene regulation, the determination of the chromatin state for each TFBS remains experimentally difficult and expensive, while computational inference from limited experimental evidence is common in the literature. For instance, CENTIPEDE [31] is probably one of the first computational methods aiming to decipher which TFBS are actually bound at certain experimental condition instead of just defining TFBS from databases such as JASPAR [19]. CENTIPEDE makes use of DNase-seq data in an unsupervised learning algorithm to infer which TFBS are in an open active state and can compare its results with experimental data. Currently, computational analysis has at its disposal several tools to process experimental data related to gene regulation from which choosing is not an easy task. Nonetheless, some collaborative projects employ reliable pipelines, e.g. the TCGA workflow [37] or the ENCODE data processing pipelines (https://github.com/ENCODE-DCC). Often, those computational tools do not provide an intuitive interface, relying entirely on command-line instructions and/or do not report figures to interpret results from such data. For example, CENTIPEDE is a R package and, therefore, requires a minimum coding expertise. Moreover, there are other tools such as Anchor, a python package [23], Mocap, a python and R hybrid package, [10], and TEPIC, a C++ program [34]. All these methods aim to determine DNA occupancy by TFs, but require expertise from users in compiling, installing dependencies, coding, and the use of the command-line interfaces.

To overcome these difficulties, we created an efficient and easy to use method, *Tool Weighted EpigenOmic Network* (Fly T-WEoN), that is able to generate *Drosophila melanogaster* context-specific Gene Regulatory Networks (GRNs). This method employs a series of filters, that once applied to a reference network, remove TF-gene regulations that are unlikely taking place according to current knowledge on the relationship between epigenetic and TFBS activation. Specificity on resulting networks is provided by the time and context for which the omic data employed by each filter was generated. Our tool is available as a Cytoscape application that provides a user-friendly and intuitive interface where researchers easily introduce their data processed with standard protocols to generate context-specific GRNs.

## 2 Methods

### 2.1 Construction of a Reference GRN

A Reference GRN is a network that contains all known regulatory interactions between gene products and genes, regardless of developmental stage, environment or cell type in an organism. To create a reference network for *D. melanogaster*, we combined TFBS information from the ENCODE data repository [13] and FlyBase [18] to then infer regulatory relationships based on distance of TFBSs to the Transcription Start Site (TSS) of each gene in the genome of the fruit fly version 6.32 (see Supp. File NetsInfo for details). To determine whether a TF regulates a gene, we chose distance thresholds between TFBSs and the TSS of each gene, so if the TFBSs falls within this distance, we assumed it regulates the respective gene. We created three reference networks with different distance thresholds, 1500, 2000, and 5000 nucleotides inspired by other approaches [6]. In the case of miRNA, genetic relationships based on experimentally determined targets from miRecords [39] and miRTarBase [11] were also retrieved and incorporated into the reference networks.

### 2.2 Filtering the reference network

In order to determine which regulatory relationships are taking place in any experimental context of interest, we defined several filters, each relying on a different type of experimental data as input. The filtering process was implemented in PERL and is the backend software of the Cytoscape [12] application developed to provide a tool with a user-friendly interface. The filtering procedure generates a time and tissue-specific GRN depending on the experimental condition in which experimental data used was generated.

Our method considers experimental information following this order for each TFBS: chromatin accessibility (DNase-seq), methylation of the DNA, histones modifications around the TFBS, the expression of each TF with known TFBSs in the reference network and miRNA quantification (see Fig. 1). First, if there is a positive signal in the TFBSs for DNA methylation, Fly T-WEoN assumes that TFs cannot bind its TFBS and the filter removes the regulation accordingly. Second, if chromatin accessibility data, e.g., DNase-seq, shows a positive signal within the chosen distance threshold used to assign a TF to the regulation of a gene, this indicates that a TF can bind the corresponding region and therefore, the edge is not removed. The next filter considers if the chromatin is in open or closed state based on histone marks experimentally associated to this process. For example, trimethylation of the Histone H3 Lys27 [42, 7] or trimethylation of the Histone H4 Lys20 [41, 4, 20] for a inactive chromatin. The effect of the histone marks considered by default in the histone marks filter are described in Table 1, and sequencing reports in BED format were used as provided in ENCODE and FlyBase. Finally, the last filter considers if the gene coding a regulator (TF or miRNA) is expressed, regulations emerging from that node are kept in the final network.

**Table 1:** Histone modifications considered in Fly T-WEoN and their default effect. Effect of the histone marks on the binding of TFs to chromatin. “+” symbols indicate marks that allow TF binding and “-” indicate non-active TFBSs.

| Modification | Effect | References |
| --- | --- | --- |
| **H3K27me3** | - | [7, 42] |
| **H3K36me2** | + | [4] |
| **H3K36me3** | + | [4, 15, 20, 7] |
| **H3K4me1** | + | [4] |
| **H3K4me2** | + | [4, 7] |
| **H3K4me3** | + | [4, 15, 7] |
| **H3K79me2** | + | [35] |
| **H3K9ac** | + | [42, 7] |
| **H3K9me2** | - | [41, 15, 32], |
| **H3K9me3** | - | [32, 42] |
| **H3S10ph** | + | [7] |
| **H4K16ac** | + | [4] |
| **H4K20me3** | - | [4] |

**Figure 1:**
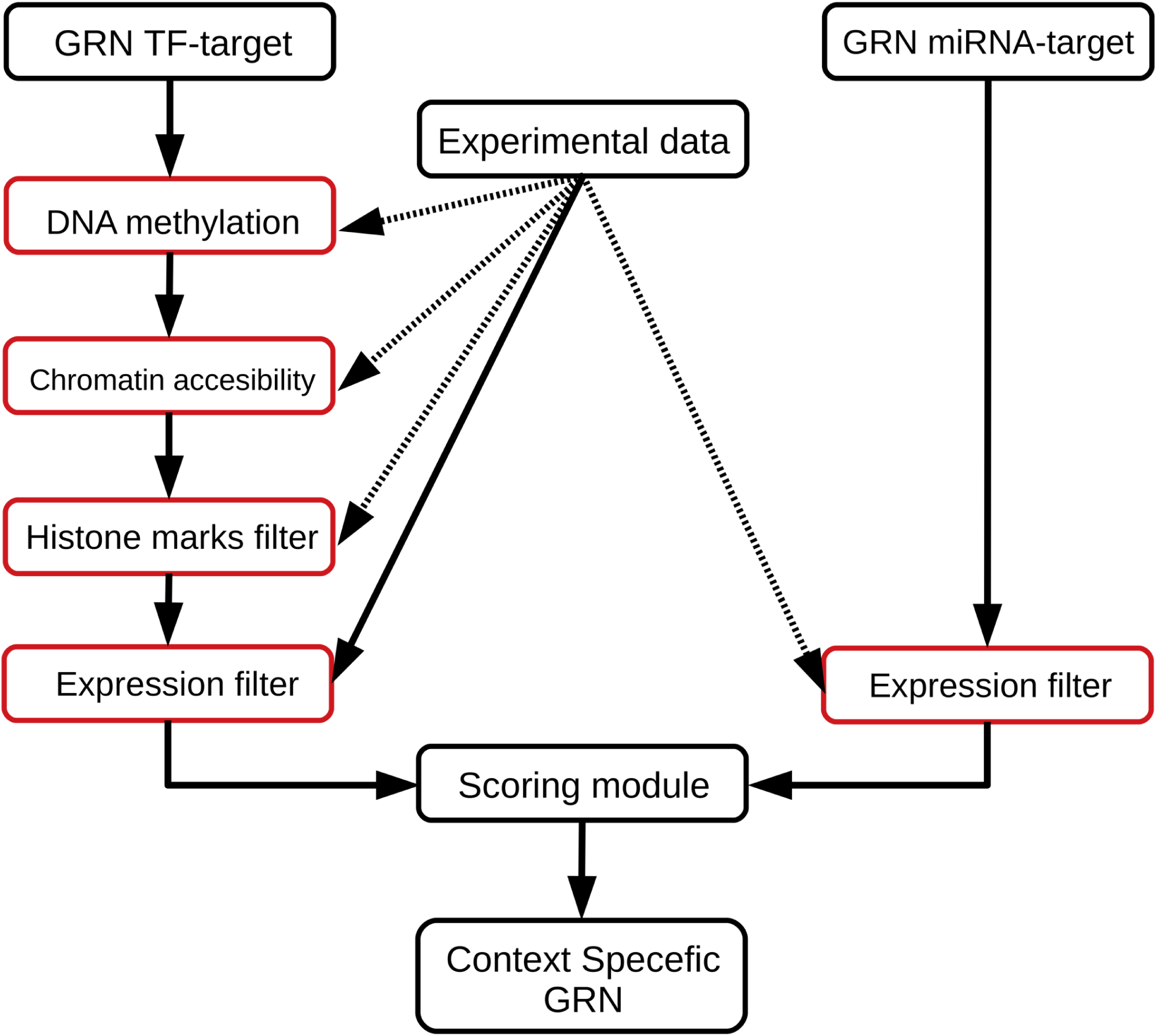
Flowchart describing Fly T-WEoN. The TF-gene reference network is filtered by DNA methylation, then chromatin accessibility of regulatory sites and third by histone marks. TF-gene and miRNA-gene networks are then filtered according to RNA-seq expression of the regulators and edges in the resulting networks are then scored accordingly to the number of filters past and provided as edge weights in the context-specific GRN.

#### 2.2.1 Scoring edges

Fly T-WEoN assigns weights to edges in the resulting network. The weight of each edge is calculated by adding a score of one for each filter that the edge passes. By default, edges have no weight, so a weight of one means the edge passed only the expression filter, a weight of two means it passed an additional filter such as a histone mark, and a weight of three indicates that the edge passed the expression filter, and for example, two different histone modifications indicated its binding site was active.

### 2.3 Validation

To assess the reliability of GRNs generated with Fly T-WEoN, we used as gold-standard a network created with all TF ChIP-seq experiments available in the ENCODE repository for the third instar larval stage or L3 of *D. melanogaster*. We chose this stage because there are ChIP-seq experiments for 32 different TF and for 10 histone marks as well as RNA-seq data. All experiments considered were carried out in equivalent conditions (see Supp. File NetsInfo for the list and IDs of experiments used). The gold-standard network was created by first removing edges from the reference network arising from genes coding for any regulator that is not among those 32 TFs, and second, by removing those edges whose TFBS was not occupied by its respective TF

#### 2.3.1 Network reliability: edges

We estimated the performance of Fly T-WEoN by considering the presence/absence of edges in the final network as a binary classification problem. In this set-up, a True Positive (TP) is defined as an edge present in the context-specific network generated after applying the filters and in the gold-standard network. Similarly, a False Negative (FN) edge is absent in the network generated by Fly T-WEoN but it is present in the gold-standard, while a False Positive (FP) edge is present in the network and absent in the gold-standard. Importantly, True Negatives (TNs) indicate edges absent in both the gold-standard and in the network created by Fly T-WEoN. Finally, once all edges are assigned to either of the three types TP, FP or FN, they were used to calculate Precision (P, eq.), Recall (R, eq.), and F1 (eq.), metrics that serve as indicators of the reliability of the context-specific networks. Each of these metrics has a value in the [0,1] range, with greater values indicating a better classification. To evaluate the effect of distance threshold we also calculated the performance metrics using the reference networks generated using the three distance thresholds 1.5, 2 and 5 Kb (See Supp. file NetsInfo). 

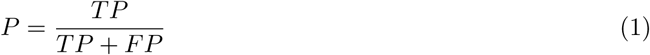

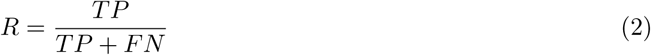

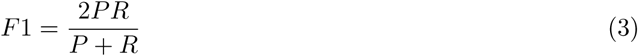

#### 2.3.2 Network reliability: local topology

GRNs are formed by combinations of graphlets, induced subgraphs that have been associated to specific functions [29]. Graphlets can be used to describe local topology of nodes in GRNs, and the presence or absence of the graphlets in which a node participate indicate functional variation for that gene in two realizations of the same network [27]. In addition, the presence or absence of graphlets in two versions of the same network can be considered as a binary classification problem, and thus, the same metrics calculated for edges indicate how similar is the local topology of each gene in the gold-standard network and in the predicted GRNs, or their overall topological similarity. We employed LoTo [26] to calculate precision, recall and the F1 metrics calculated for the presence/absence of graphlets in every pairwise network comparison. If these metrics only consider graphlets in which the same gene participates, they serve to indicate variations in the local topology of that node. Whereas, if the metrics are calculated for all graphlets in the networks, they serve to indicate global topological similarity between the two networks.

### 2.4 Fruit fly early embryo development

To demonstrate the utility of Fly T-WEoN, we generated networks for six different stages of early embryo development in fruit fly (*D. melanogaster*). We employed RNA-seq experiments and histone marks data downloaded from different databases such as modENCODE and modMine projects [9, 13], and the FlyBase database [18] (see Supp. file NetsInfo for a detailed description of the data used).We downloaded the annotation of the *D. melanogaster* reference genome version 6.32 to process all sequencing experiments. Experiments already mapped to a different version of the reference genome were re-processed or converted using the FlyBase Sequence Coordinates Converter [18]. We employed these data to create context-specific networks for different time points of early development of *D. melanogaster*. The default

1.5kb reference network that is included in Fly T-WEoN was used for this example. This reference network comprises 15576 genes (87% of the total annotated genes of *D. melanogaster*). Six time-specific networks were created with Fly T-WEoN encompassing the fly embryonic development (0-24h) in time steps of 4 hours (0-4h, 4-8h, 8-12h, 12-16h, 16-20h, and 20-24h), using the available data of histone modifications and RNA-seq.

Finally, we compared each of these networks with the network created for the consecutive time interval using LoTo [26] to calculate overall network similarity and to identify genes whose local topology changed during embryo development according to the F1 calculated for all graphlets in which they participate. For each comparison, we separated nodes by their type (TFs, non-TF protein coding genes and non-coding genes) into four F1 intervals [0-0.5), [0.5-0.7), [0.7-0.9) and [0.9-1.0). For those coding genes that are not TFs in each of these intervals we determined the statistical over-representation of GO-Slim Molecular Process terms with PANTHER using the Fisher’s exact test with the Bonferroni correction [28].

## 3 Results

### 3.1 Method validation

We employed the L3 context-specific GRN described in the Methods section to estimate the reliability of the networks generated by our approach. The gold-standard network was made with 32 different TFs and their binding sites determined by ChIP-seq experiments in equivalent experimental conditions. The reference networks made at 1.5, 2 and 5 kb thresholds are described in table 2. Not surprisingly, larger distance thresholds include more TF-gene interactions for genes to which we can not assign regulators otherwise, and thus, networks built using greater thresholds contain more nodes.

**Table 2:** Gold standard Networks used to validate Fly T-WEoN. Networks made with the 32 TFs at different distance thresholds between TFBSs and the TSS of each gene. Number of different genes and edges present in each of the networks made by assigning a TF to the regulation of a gene if the TF is bound within the distance and the gene TSS.

| Threshold (Kb) | Genes | Edges |
| --- | --- | --- |
| 1.5 | 10096 | 82919 |
| 2 | 10620 | 89880 |
| 5 | 12822 | 127222 |

#### 3.1.1 Network similarity: edges

Using the L3 example, the lowest score of edges in the predicted networks is two and the highest eleven. This is due to the number of Fly T-WEoN filtering steps applied, so a score of two implies that the TF from which an edge is originated is expressed and there is at least a single histone modification supporting its existence. Scores of three, and above, mean that there is at least two types of histone modification indicatingthat the link exists.

As shown in Table 3 for a threshold of 1.5kb, Fly T-WEoN generates networks with very high similarity to the gold-standard network in our benchmarking. Starting with edges of score two or greater, the network generated by Fly T-WEoN contains 97.8% of the edges of the gold-standard network (R = 0.978), decreasing the recall as the edges score increases. Also, the F1 value follows the same trend, it displays its highest value using this score (F1 = 0.884) and decreases as the minimum score for the edges increases. Moreover, the precision follows a different tendency, with its highest value with score *≥* 6 (P = 0.810). The worst performance is obtained with a score of eleven, the maximum, with which Fly T-WEoN recovers 0.1% of the edges of the gold-standard network (R = 0.001, P = 0.646 and F1 = 0.001), indicating low similarity between edges present in the predicted networks and the gold-standard.

**Table 3:** Reliability of L3 gene regulatory networks: single edges. Performance of Fly T-WEoN measured by its ability to recover edges present in the gold-standard network for different scores. The table displays the number of True Positive edges (TP), edges in the gold-standard network also present in the predicted network; False positive edges (FP) or present in the predicted network but absent in the gold-standard network; and False Negative edges, those edges that are are only present in the gold-standard network and are not present in the predicted network. TP, FP and FN edges were used to calculate Precision (P), Recall (R), and F1 (bold numbers indicate their highest values).

| Score | TP | FP | FN | R | P | F1 |
| --- | --- | --- | --- | --- | --- | --- |
| 2 | 81094 | 19475 | 1825 | **0.978** | 0.806 | **0.884** |
| 3 | 78807 | 18846 | 4112 | 0.950 | 0.807 | 0.873 |
| 4 | 76017 | 17998 | 6902 | 0.917 | 0.809 | 0.859 |
| 5 | 72802 | 17109 | 10117 | 0.878 | 0.810 | 0.843 |
| 6 | 68848 | 16147 | 14071 | 0.830 | **0.810** | 0.820 |
| 7 | 61477 | 14874 | 21442 | 0.741 | 0.805 | 0.772 |
| 8 | 50071 | 12908 | 32848 | 0.604 | 0.795 | 0.686 |
| 9 | 5666 | 2225 | 77253 | 0.068 | 0.718 | 0.125 |
| 10 | 1512 | 664 | 81407 | 0.018 | 0.695 | 0.036 |
| 11 | 62 | 34 | 82857 | 0.001 | 0.646 | 0.001 |

#### 3.1.2 Global topological similarity calculated with graphlets

The trend for graphlet based results is similar to that based on single edges, shown in Table 4. Using a minimum score of two, Fly T-WEoN is able to recover 95.7% of the graphlets found in the Gold Standard Network (R = 0.957), but it tends to over predict graphlets as indicated by the much lower precision (P = 0.662). Also, the F1 value had its greatest value with a score of at least two (F1 = 0.782), indicating again high similarity between the predicted and gold-standard networks. The highest value of precision was obtained using at minimum score of five (P = 0.665), this indicating again that networks obtained by Fly T-WEoN contain more graphlets than gold-standard networks, even at the maximum precision. The lowest values for the performance metrics were obtained using weights *≥* 10, with predicted networks recovering 0% of the graphlets present in the gold-standard network.

**Table 4:**
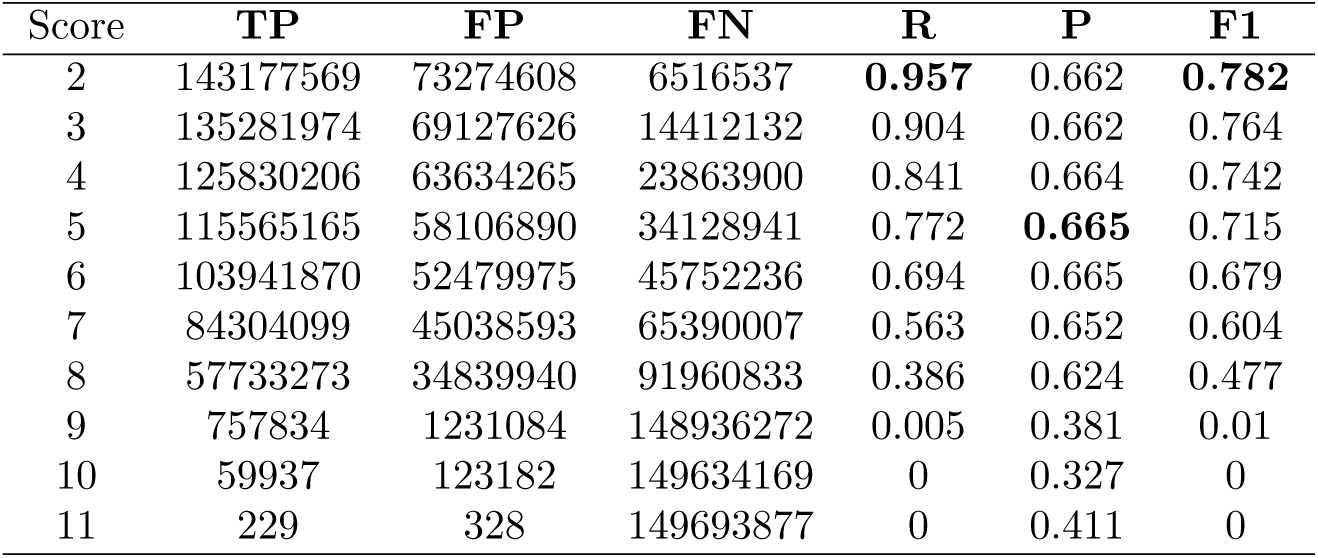
Reliability of L3 gene regulatory networks: graphlets. Performance of Fly T-WEoN measured by its ability to recover graphlets present in the gold-standard network for different edge scores. The table displays the number of True Positive graphlets (TP), graphlets present in the gold-standard network also found in the predicted network; False positive graphlets (FP), present in the predicted network but absent in the gold-standard network; and False Negative graphlets, those that are are only present in the gold-standard network and were not present in the predicted network. TP, FP and FN graphlets were used to calculate Precision (P), Recall (R), and F1 (bold numbers indicate their highest values).

### 3.2 An example case: Fruit fly early development

#### 3.2.1 Network sizes

Six time-specific networks were created with Fly T-WEoN encompassing the fly embryonic development (0-24h) in consecutive time ranges of 4 hours (0-4h, 4-8h, 8-12h, 12-16h, 16-20h, and 20-24h). These networks were made using available data of histone modifications and RNA-seq. These networks have different numbers of edges, graphlets, regulatory nodes (TFs) and total number of genes, as shown in Table 5. The largest network belongs to the 16-20h time range, with the largest numbers for nodes, total connections, and regulatory nodes (10993, 928599, and 345 respectively). The smallest network is the network for the 0-4h time range, which has the lowest number of total connections, and regulatory nodes (718583 and 305 respectively), while the network for time range 4-8h has the lowest number of nodes (7886).

**Table 5:** Characterization of embryo development networks. The table shows the number of edges and regulatory nodes for each of the networks created for the 6 time intervals during early development of the fruit fly. Regulatory nodes indicate the number of TFs in each network and the total number of genes and edges in the networks are also displayed. These networks were obtained by removing unlikely edges from a reference networks were TFBSs located at most at 1.5kb upstream the TSS are used to assign the TFs that bind to that TFBS to the regulation of each gene.

| Graphlet type | 0-4h | 4-8h | 8-12h | 12-16h | 16-20h | 20-24h |
| --- | --- | --- | --- | --- | --- | --- |
| **Total edges** | 718583 | 733863 | 803613 | 888537 | 928599 | 840567 |
| **TF nodes** | 305 | 324 | 340 | 335 | 345 | 339 |
| **Total nodes** | 8811 | 7886 | 8554 | 10528 | 10993 | 11146 |

#### 3.2.2 Network Comparisons

We compared each network with the network representing the next time interval obtaining F1 values greater than 0.85 (Table 6). These results indicate that despite changes, most of the regulatory network remains unaltered between time lapses. Thus, indicating that relatively small changes in the network account for all stages of early embryo development.

**Table 6:** Comparisons of embryo development networks using gaphlets. The table displays the number of True Positive graphlets (TP), graphlets in the first network (that belonging to the earlier time interval) that are present in the later network; False positive graphlets (FP) are those present in the later network but absent in the earlier one; and False Negative graphlets (FN), those that are are only present in the earlier network and not in the later network. TP, FP and FN graphlets were used to calculate Precision (P), Recall (R), and F1 metrics.

| Comparison | TP | FP | FN | R | P | F1 |
| --- | --- | --- | --- | --- | --- | --- |
| 0-4h – 4-8h | 960856575 | 154840150 | 170821683 | 0.849 | 0.861 | 0.855 |
| 4-8h – 8-12h | 1015103337 | 237272519 | 100593375 | 0.910 | 0.811 | 0.857 |
| 8-12h – 12-16h | 1162138649 | 358797508 | 90237194 | 0.928 | 0.764 | 0.838 |
| 12-16h – 16-20h | 1347047992 | 264431780 | 173888152 | 0.886 | 0.836 | 0.860 |
| 16-20h – 20-24h | 1297280051 | 74185782 | 314199708 | 0.805 | 0.946 | 0.870 |

We also analyzed the F1 values by types of genes, TF and non-TF coding, and non-coding genes (Table 7). Without considering gene type (all genes), most of them are in the F1 ranges with less topological variation ([0.7, 0.9) and [0.9,1.0]), evidencing that, as happened with global topology, the local topology of a majority of genes remains unaltered between consecutive time lapses. The same trend is displayed by the TF-coding genes, with most of them in the range [0.7,0.9). With respect to non-TF coding genes, again most of them fall into F1 intervals ranges with less topological variation ([0.7,0.9) and [0.9,1.0]). Notably, there are large proportions of ncRNA coding genes in the range that displays larger topological variations, hinting they play a relevant role in the developmental stages depicted by the networks. Detailed information on which genes show greater variation on their local topology and the GRN for each time point can be found in the Supp. Material of the article (file LoTo Embryo and EmbryoNetworks respectively).

**Table 7:** Total number of genes by type and F1 interval in each of the comparisons of embryo development consecutive networks using gaphlets. The table displays the number of genes in each of the four F1 intervals, [0.0,0.5), [0.5,0.7), [0.7,0.9) and [0.9,1.0] in each of the five comparisons performed between GRNs depicting gene regulation at each time lapse. F1 one values closer to 0 indicate larger local topological variation, while closer to 1 indicate fewer variations in the graphlets in which a gene participates.

|  | Comparison | 0-4h – 4-8h | 4-8h – 8-12h | 8-12h – 12-16h | 12-16h – 16-20h | 16-20h – 20-24h |
| --- | --- | --- | --- | --- | --- | --- |
| All genes | [0.0,0.5) | 1459 | 1398 | 2595 | 2962 | 2511 |
|  | [0.5,0.7) | 187 | 178 | 215 | 319 | 397 |
|  | [0.7,0.9) | 3192 | 1732 | 2023 | 1766 | 1546 |
|  | [0.9,1.0] | 4097 | 5445 | 5820 | 6857 | 7513 |
| TFs | [0.0,0.5) | 27 | 25 | 10 | 5 | 10 |
|  | [0.5,0.7) | 7 | 9 | 12 | 7 | 4 |
|  | [0.7,0.9) | 212 | 260 | 302 | 273 | 265 |
|  | [0.9,1.0] | 70 | 47 | 16 | 53 | 66 |
| Coding genes | [0.0,0.5) | 953 | 922 | 1787 | 2114 | 1802 |
|  | [0.5,0.7) | 136 | 136 | 151 | 236 | 292 |
|  | [0.7,0.9) | 2602 | 1242 | 1382 | 1126 | 973 |
|  | [0.9,1.0] | 3598 | 4815 | 5077 | 5925 | 6527 |
| Non-coding genes | [0.0,0.5) | 479 | 451 | 798 | 843 | 699 |
|  | [0.5,0.7) | 42 | 33 | 52 | 76 | 101 |
|  | [0.7,0.9) | 378 | 230 | 339 | 367 | 308 |
|  | [0.9,1.0] | 429 | 583 | 727 | 879 | 920 |

#### 3.2.3 Functional analysis of genes with altered local topology

After performing comparisons of networks representing consecutive developmental stages, we analyzed the function of genes with altered local topology. To do so, we employed the statistical enrichment of GO-Slim Biological Process terms with PANTHER [38] for genes in each of F1 ranges previously defined. Focusing on the analysis of genes with F1 in the [0-0.5), the enrichment test denoted several GO terms that are known to be involved in embryonic development (Table 8). For example, we found enriched GO terms “developmental process” and “anatomical structure development” in genes in the lowest F1 range in the comparisons spanning the first 12 hours (0-4h-4-8h and 4-8h-8-12h). In the comparisons spanning the last 12 hours, we found enriched functional terms related to metabolism and metabolite transport processes such as “glutathione metabolic process”, “transmembrane transport”, and “aminoglycan metabolic process”. Genes in intervals with moderate topological variation (F1 range [0.5-0.7), see Supp Material GO file) showed enrichment in GO terms related to defense response, metabolic, and developmental process. For the comparison of 4-8h and 8-12h networks, genes in this F1 range, enriched terms were “animal organ development”, “cytoplasmic translation”, and “cell development”. In the case of the comparison 8-12h–12-16h, enriched terms associated to cell signaling, GO terms “signaling” and “cell communication”. Finally, for the comparison of 12-16h–16-20h, overrepresented terms were related with cell structure and cell cycle, GO Slim terms such as “establishment of spindle orientation” and “cell cycle”. In the case of the comparison 16-20h – 20-24 no GO term was significantly enriched.

**Table 8:** GO Slim Biological Process terms associated to genes with the largest topological variation. The table displays the GO Slim Biological Process obtained with PANTHER for genes with F1 values in the range [0.0 - 0.5). The Fold Enrichment value indicate the rate between the percentage of genes with the annotation and the percentage of genes with the same annotation in whole genome. If it is greater than 1, it indicates that the category is overrepresented in the data. These results were filtered by a p-value threshold of 0.01.

| Comparison | GO Term | GO ID | Fold Enrichment | p-value |
| --- | --- | --- | --- | --- |
| 0-4h – 4-8h | cell differentiation | GO:0030154 | 2.29 | 1.14e-04 |
|  | developmental process | GO:0032502 | 2.02 | 1.49e-02 |
|  | cellular developmental process | GO:0048869 | 2.15 | 3.12e-04 |
|  | sulfur compound metabolic process | GO:0006790 | 2.53 | 1.06e-03 |
|  | anatomical structure development | GO:0048856 | 1.92 | 1.41e-03 |
|  | cellular modified amino acid metabolic process | GO:0006575 | 2.94 | 1.85e-03 |
|  | glutathione metabolic process | GO:0006749 | 3.52 | 2.40e-03 |
|  | cell fate commitment | GO:0045165 | 4.82 | 3.54e-03 |
|  | neurogenesis | GO:0022008 | 2.38 | 4.14e-03 |
|  | generation of neurons | GO:0048699 | 2.38 | 5.81e-03 |
|  | multicellular organismal process | GO:0032501 | 1.58 | 9.06e-03 |
| 4-8h – 8-12h | developmental process | GO:0032502 | 1.87 | 1.15e-03 |
|  | cell differentiation | GO:0030154 | 2.04 | 2.07e-03 |
|  | chaperone-mediated protein folding | GO:0061077 | 4.18 | 3.16e-03 |
|  | cellular developmental process | GO:0048869 | 1.91 | 4.46e-03 |
|  | anatomical structure development | GO:0048856 | 1.79 | 6.33e-03 |
| 8-12h – 12-16h | cellular modified amino acid metabolic process | GO:0006575 | 4.18 | 2.61e-09 |
|  | glutathione metabolic process | GO:0006749 | 5.00 | 2.04e-08 |
|  | cofactor metabolic process | GO:0051186 | 2.24 | 3.25e-05 |
|  | sulfur compound metabolic process | GO:0006790 | 2.32 | 1.10e-04 |
|  | response to drug | GO:0042493 | 2.92 | 1.39e-03 |
|  | organic acid metabolic process | GO:0006082 | 1.66 | 2.81e-03 |
|  | small molecule metabolic process | GO:0044281 | 1.46 | 3.85e-03 |
|  | transmembrane transport | GO:0055085 | 1.55 | 6.24e-03 |
|  | carboxylic acid metabolic process | GO:0019752 | 1.62 | 6.36e-03 |
|  | organic anion transport | GO:0015711 | 2.05 | 6.50e-03 |
|  | oxoacid metabolic process | GO:0043436 | 1.60 | 6.65e-03 |
|  | aminoglycan metabolic process | GO:0006022 | 3.43 | 6.94e-03 |
|  | anion transport | GO:0006820 | 1.84 | 8.37e-03 |
|  | defense response | GO:0006952 | 3.25 | 8.89e-03 |
| 12-16h – 16-20h | aminoglycan metabolic process | GO:0006022 | 3.98 | 7.78e-04 |
|  | amino sugar catabolic process | GO:0046348 | 3.62 | 2.28e-03 |
|  | chemical synaptic transmission | GO:0007268 | 2.09 | 2.73e-03 |
|  | anterograde trans-synaptic signaling | GO:0098916 | 2.09 | 2.73e-03 |
|  | response to drug | GO:0042493 | 2.64 | 3.05e-03 |
|  | synaptic signaling | GO:0099536 | 2.06 | 4.36e-03 |
|  | trans-synaptic signaling | GO:0099537 | 2.06 | 4.36e-03 |
|  | multicellular organismal process | GO:0032501 | 1.41 | 7.76e-03 |
| 16-20h – 20-24h | aminoglycan metabolic process | GO:0006022 | 4.25 | 7.65e-04 |
|  | amino sugar catabolic process | GO:0046348 | 4.25 | 7.65e-04 |
|  | drug metabolic process | GO:0017144 | 1.85 | 5.51e-03 |

#### 3.2.4 Subnetworks of nodes showing largest topological variations at early stages

To further investigate the application of our approach to the early embryo development example, we created subnetworks made of only those nodes that have F1 in the [0.0,0.5) range for each comparison. We then compared subnetworks depicting consecutive stages using LoTo. As an example we show the comparison of the two earlier stages (0-4h – 4-8h) in figures 2 and 3 the results of the comparison showing only TFs. All these subnetworks can be found as a Cytoscape session in the Supp. Material of the article.

**Figure 2:**
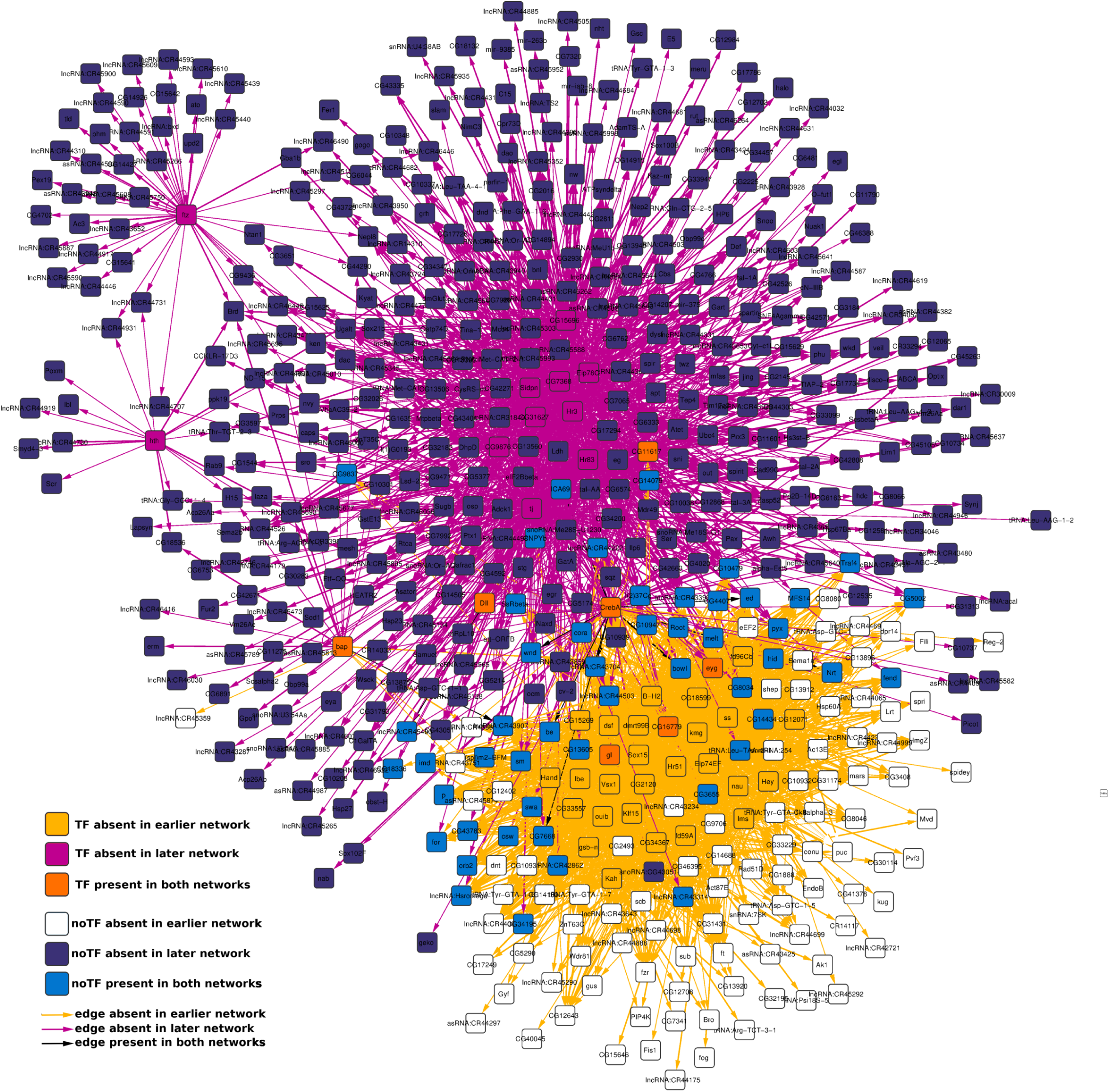
Comparison of subnetworks composed by all those genes showing larger local variation in the 0-4h – 4-8h comparison. The network shown is formed by 594 nodes (44 TFs) and 3107 edges colored accordingly to their existence in the earlier network, in the later network or in both.

**Figure 3:**
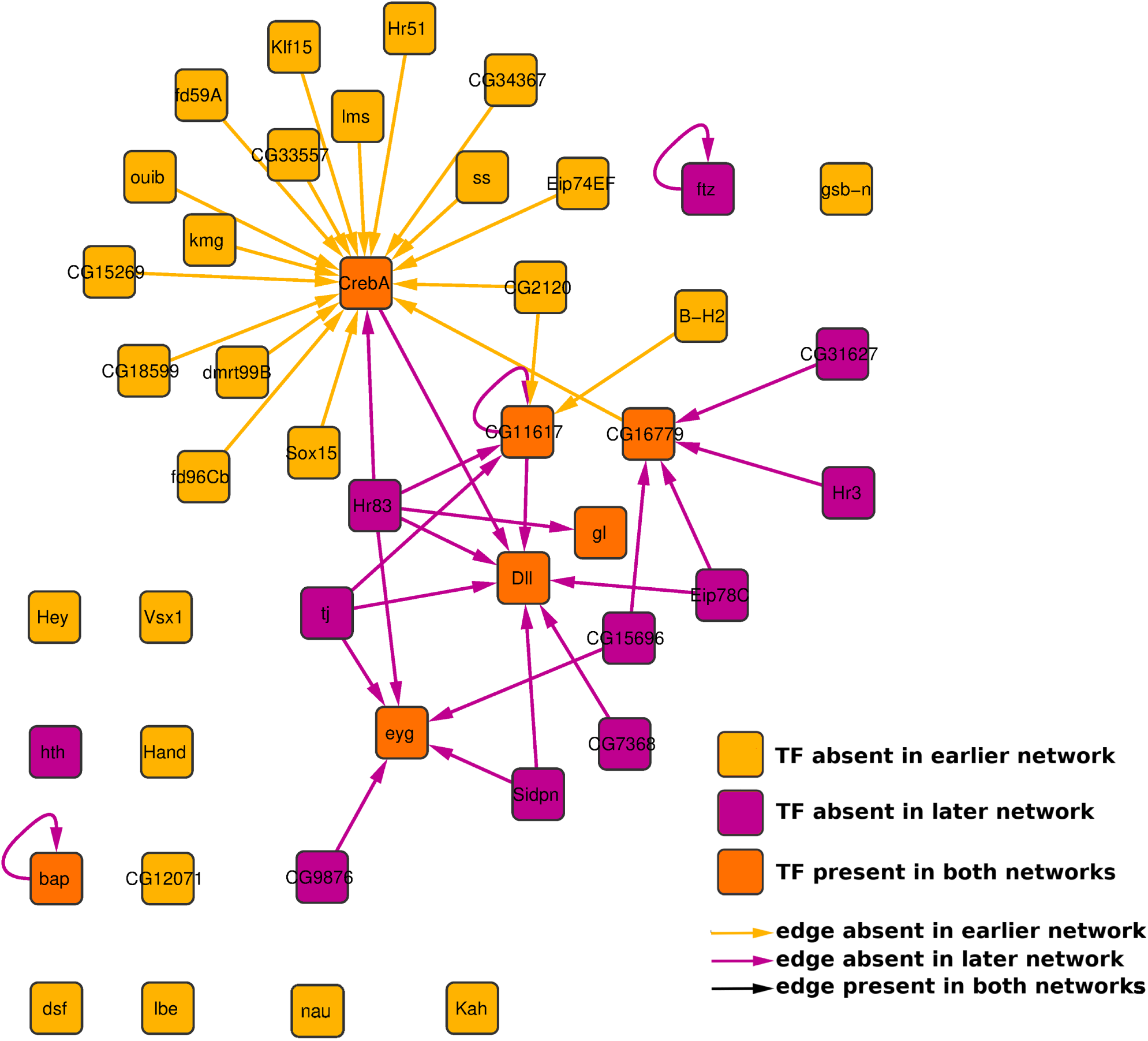
Comparison of subnetworks composed by TF coding genes showing larger local variation in the 0-4h – 4-8h comparison. The network shown is formed by 44 TFs and 42 edges colored accordingly to their existence in the earlier network, in the later network or in both.

## 4 Discussion

Inference of gene regulation relationships from genomic data is a particularly hard and costly task. This is due to the use of high quality antibodies to determine the bound state of TFs to the open chromatin. Even with aid of computational tools, the determination of gene regulations is an open problem contributed by a gap knowledge of how TFs and other regulators of gene expression work, and by a general lack of genomic data suitable for the prediction of such regulations. The inexpensive RNA-seq and chromatin accessibility through footprint sequencing are commonly used to infer condition-specific networks, but these still require corroboration that again, is usually made with comparisons to ChIP-seq experiments of each TF. However, TF ChIP-seq experiments are unavailable for most conditions of model organisms, including even those that have been deeply studied. Importantly, the number of ChIP-seq used to determine histone modifications are increasing in data repositories, and given the relationship between histone marks and TF binding in chromatin, we created Fly T-WEoN to generate context-specific GRNs in *D. melanogaster*.

With the proposed methodology, we built condition-specific GRNs for the L3 developmental stage and used them to validate our methodology based on the concatenation of simple filters. We employed ChIP-seq for ten different histone marks and 32 different TFs to build gold-standard networks to then compare them Fly T-WEoN networks. Each of the filters in our tool uses current knowledge on the known relationship that exists between epigenetic marks and TF activity. We observed that even if the filtering approach may seem to be too simple it still recovers correctly most of the edges found in the Gold Standard Network we made for that stage. Furthermore, Fly T-WEoN applies a weight system on edges, increasing these weights accordingly to how many filters did each edge past. Our results show that, at least in our test, using a weight of *≥* 2 produces the most reliable GRNs. This weight means that at least one histone mark and the expression of the TF agrees with the existence of each edge. The worse performance shown with greater weights can be explained due to that by increasing the weight value, the number of edges and graphlets in the networks decrease. However, using only edges with greater weights decreases the reliability of the edges (tables 3 and 4). Which suggests that the known effect of different epigenetic marks is contradictory, and thus, our simple filtering approach fails to gather the complexity of the epigenetic code.

To highlight the differences and similarities between Fly T-WEoN and other approaches, we report a brief comparison between Fly T-WEoN and other four methods in Table 9. The other methods used for the comparison were CENTIPEDE [31], Anchor [23], TEPIC [34], and Mocap [10]. It is important to stress that none of these methods was designed or even tested for *D. melanogaster*, and thus a quantitative comparison is not straightforward. Given the heterogeneous data employed by these methods, the absence of actual context-specific GRNs, and the lack of specific tools for *D. melanogaster*, it is not possible to perform quantitative comparisons between them, and thus, only qualitative comparisons are possible. Our comparison (see Table 9) highlights the main characteristics of Fly T-WEoN, i.e., the intuitive way to use Fly T-WEoN and the integration of its results in Cytoscape, when compared with the other four approaches. It is very important to highlight that these tools use different types of data (see Table 9) in dissimilar context to those used by Fly T-WEoN. This makes even more difficult to make a quantitative performance comparison between them. Also, there is no-context-specific data available for all data types used by Fly T-WEoN (DNAse, RNAseq, DNA methylation and TFs ChIP-seq), which does not allow for a full comparison.

**Table 9:**
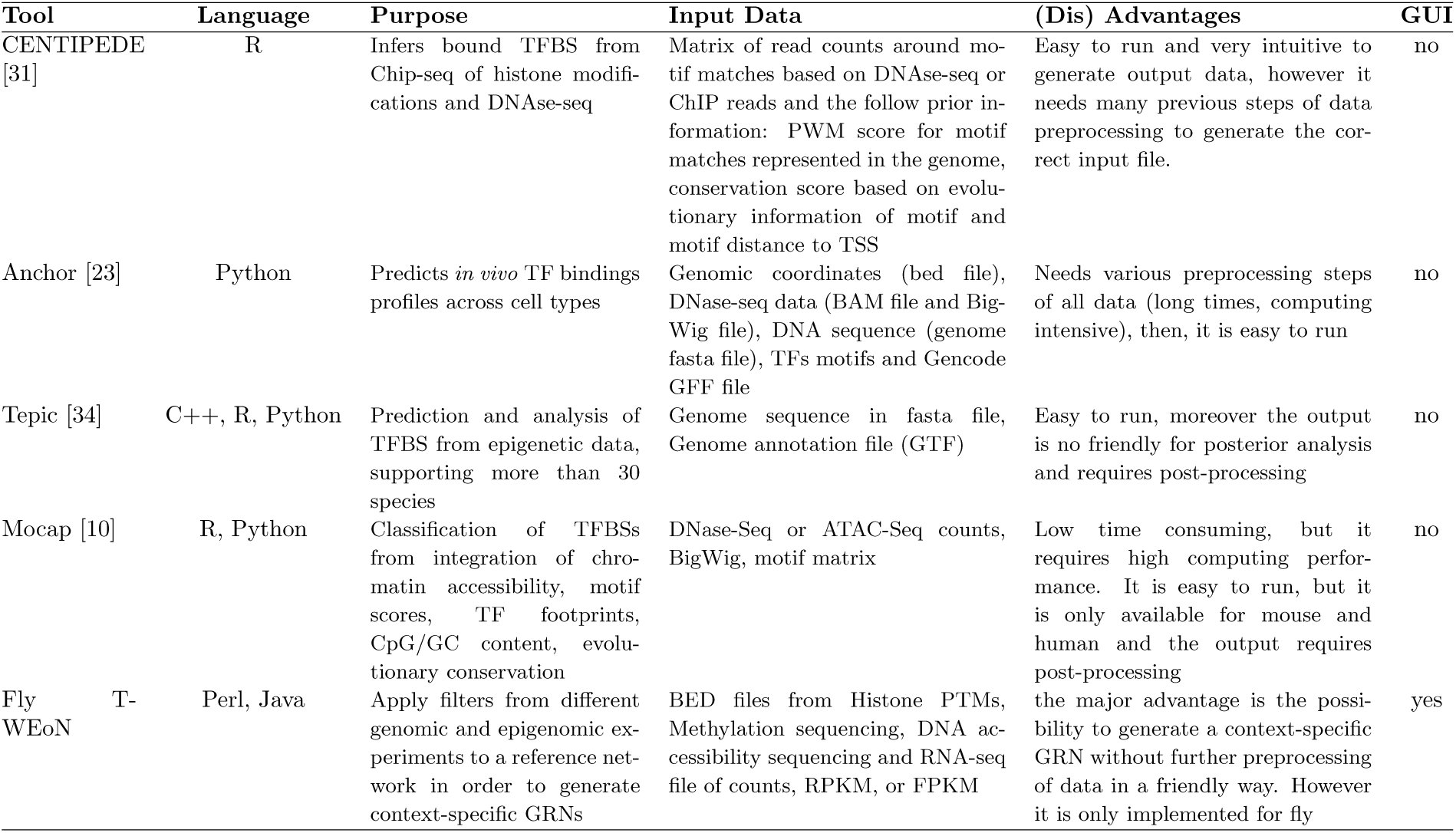
Qualitative comparison of different methods and Fly T-WEoN. The table indicates for each tool the language used in its implementation, its purpose, its advantages and disadvantages and general user-friendliness.

Regarding the example of embryonic development of *D. melanogaster*, we created 6 different networks, each depicting transcriptional control by TFs for each of the four hours intervals of the first 24 hour of a fly embryo. We opted for this condition and time intervals because these were the conditions for which there is more epigenetic and transcriptional data at modENCODE and GEO datasets. Importantly, the stages represented by our networks are when cells and tissues in *D. melanogaster* are more homogeneous, and thus, all omic data employed is deemed to be more significant. When comparing these GRNs with LoTo [26], we observed that the networks increase the number of nodes and connections as development progresses. This may indicate that in later stages of development transcriptional regulation becomes a more complex process that involves a grater number of TFs in greater number of cell fates and tissues. Comparisons of overall similarity between networks representing consecutive time intervals showed that the largest variation takes place between 8-12h and 12-16h networks and that the overall topology of the networks changes less is in the last transition between developmental stages included, i.e. in the comparison 16-20h and 20-24h networks.

With respect to variations in the local topology of single nodes determined by F1 calculated for the presence/absence of graphlets, most genes had small variations in all comparisons 7, a trend observed for all gene types in the networks (TFs, protein coding and ncRNAs). The only exception is ncRNA coding genes, mainly lncRNAs, which are almost as numerous in the F1 range that indicates largest topological variation as in the range depicting the lowest variation. These findings agree with previous knowledge on the role played by lncRNA in *D. melanogaster* development [24]. Regarding our observation of relatively few TFs displaying large variations in their local topology, and that those with larger changes (lowest F1) are densely linked between them, these findings agree with the concept of clusters of master regulators [14]. In this concept, a small cluster of highly interconnected TFs are the master regulators controlling the other regulators whose function is to act as efectors or “fine-tuners” of the orders given by the master regulators. In our example, regarding the master regulator concept, the “fine-tuners” would regulatory nodes found in the F1 ranges with higher values that are linked to the master regulators and to many other genes that do not code for regulators. Nonetheless, it should also be considered that specially at the earlier stages of embryo, there are many TFs that are inherited from the mother [1], and given that our approach uses as approximation for TF activity the expression of their coding genes, maternal TFs are disregarded. The fact of observing an increasing number of nodes as the networks depict later stages also agrees with known facts regarding developments, as tissues and specialized cells appear, both regulators and non regulator genes tend to perform more specialized functions [1]. Our functional analysis also corroborates this (see Table 8), more general functions related at the earlier stages and more specialized functions as development progresses, validating again the networks generated with Fly T-WEoN.

## 5 Conclusion

Here, we demonstrated the reliability of our tool, Fly T-WEoN, with results indicating that most of the regulatory events depicted by edges in its resulting networks are likely taking place. In addition to this validation, and given the current lack of tools that integrate epigenetic data for the construction of GRNs in *D. melanogaster*, we also provided a qualitative comparison with other approaches, helping in this way to stress out the usability of our method. The minimum input required by Fly T-WEoN is a quantification of the expression of genes, but the results we show here prove how the quality of the network improves by using other epigenetic data or quantification of miRNAs.

We finally demonstrated through a case study the usefulness of genomic data to filter out known regulations from a reference network and make context-specific gene regulatory networks where functions of genes with varying regulation correlate with the development stage. Moreover, we developed a Cytoscape app for Fly T-WEoN that serves as frontend for the presented method, allowing users to create and visualize context-specific GRNs from their processed RNA-seq, DNase-seq, bisulphite-seq, and ChIP-seq datasets or data obtained from public databases. We expect to further develop a backend software harnessing Machine Learning algorithms that would allow final users to predict gene expression from minimal and cheap genomic data, and extend the current method from fruit fly to other model organisms, specially human.

Fly T-WEoN can be obtained free of charge here https://weon.readthedocs.io. Supp. Material files con be accessed here https://figshare.com/projects/WEoN_FlyT/76983.

## Supporting information

https://figshare.com/projects/WEoN_FlyT/76983

## Acknowledgements

We acknowledge the help received from Dr. Inti Pedroso for his patience and useful discussions and Dr Yesid Cuesta for his constructive review of the manuscript.

